# Artificial intelligence predicts the immunogenic landscape of SARS-CoV-2: toward universal blueprints for vaccine designs

**DOI:** 10.1101/2020.04.21.052084

**Authors:** Brandon Malone, Boris Simovski, Clément Moliné, Jun Cheng, Marius Gheorghe, Hugues Fontenelle, Ioannis Vardaxis, Simen Tennøe, Jenny-Ann Malmberg, Richard Stratford, Trevor Clancy

**Affiliations:** NEC OncoImmunity AS, Oslo Cancer Cluster, Ullernchausseen 64/66, 0379 Oslo, Norway; NEC Laboratories Europe GmbH Kurfuersten-Anlage 36, 69115 Heidelberg, Germany

## Abstract

The global population is at present suffering from a pandemic of Coronavirus disease 2019 (COVID-19), caused by the novel coronavirus Severe Acute Respiratory Syndrome Coronavirus 2 (SARS-CoV-2). The goals of this study were to use artificial intelligence (AI) to predict blueprints for designing universal vaccines against SARS-CoV-2, that contain a sufficiently broad repertoire of T-cell epitopes capable of providing coverage and protection across the global population. To help achieve these aims, we profiled the entire SARS-CoV-2 proteome across the most frequent 100 HLA-A, HLA-B and HLA-DR alleles in the human population, using host-infected cell surface antigen presentation and immunogenicity predictors from the *NEC Immune Profiler* suite of tools, and generated comprehensive epitope maps. We then used these epitope maps as input for a Monte Carlo simulation designed to identify statistically significant “epitope hotspot” regions in the virus that are most likely to be immunogenic across a broad spectrum of HLA types. We then removed epitope hotspots that shared significant homology with proteins in the human proteome to reduce the chance of inducing off-target autoimmune responses. We also analyzed the antigen presentation and immunogenic landscape of all the nonsynonymous mutations across 3400 different sequences of the virus, to identify a trend whereby SARS-COV-2 mutations are predicted to have reduced potential to be presented by host-infected cells, and consequently detected by the host immune system. A sequence conservation analysis then removed epitope hotspots that occurred in less-conserved regions of the viral proteome. Finally, we used a database of the HLA genotypes of approximately 22 000 individuals to develop a “digital twin” type simulation to model how effective different combinations of hotspots would work in a diverse human population, and used the approach to identify an optimal constellation of epitopes hotspots that could provide maximum coverage in the global population. By combining the antigen presentation to the infected-host cell surface and immunogenicity predictions of the *NEC Immune Profiler* with a robust Monte Carlo and digital twin simulation, we have managed to profile the entire SARS-CoV-2 proteome and identify a subset of epitope hotspots that could be harnessed in a vaccine formulation to provide a broad coverage across the global population.

## Introduction

The outbreak of Coronavirus disease 2019 (COVID-19) and its rapid worldwide transmission resulted in the World Health Organization (WHO) declaring COVID-19 as a pandemic and global health emergency. COVID-19 is caused by the novel coronavirus Severe Acute Respiratory Syndrome Coronavirus 2 (SARS-CoV-2) [1]. Like all coronaviridae, SARS-CoV-2 is a positive-sense RNA viruse encapsulated by an envelope, and characterized by an exposed spike glycoprotein (S-protein) that is projected from the viral surface and comprises a large RNA genome [2]. Although the main structural proteins on coronaviridae, such as the S-protein, are reasonably well studied, many of the other proteins are less well characterized. Correcting this gap may be important to improve the design of therapeutic interventions [3]. This particular gap in knowledge is very relevant from the perspective of finding immunogenic targets across the entire virus proteome, in order to guide the design of effective vaccines. The SARS-CoV-2 virus is closely related in sequence identity and receptor binding to SARS-CoV [4, 5], and therefore it has been purported that one may borrow from this similarity to validate targets in potential vaccines [6, 7]. Much of the emphasis on coronaviridae vaccines to date has focused on antibody responses against the S-protein, which is the most “antibody exposed” structural protein in the virus. Although demonstrated to be effective with short-lived responses in a mouse study [8], the immune response against the S-protein of SARS-CoV is associated with low neutralizing antibody titers and short-lived memory B cell responses in recovered patients [9, 10]. Additionally, potential harmful effects of vaccines based on the antibody response to S-protein in SARS-CoV have raised possible safety concerns regarding this approach. For example, in macaque models, it was observed that anti-S-protein antibodies caused severe acute lung injury [11], and sera from SARS-CoV patients also revealed that elevated anti-S-protein antibodies were observed in those patients that succumbed to the infection[11]. When considering antibody responses to the S protein, it is also important to consider the possibility that antibody-dependent enhancement (ADE) may occur, whereby antibodies facilitate viral entry into host cells and enhance the infection of the virus[12]. It has already been demonstrated that neutralizing antibodies bound to the S protein of coronaviridae trigger a conformational change that facilitates viral entry into host cells[13]. Considering the potential for ADE, the reported short-lived antibody response [9, 10, 14], and the documented pathological consequence of S-protein specific antibodies in certain animal models, it is worth considering alternative strategies for vaccine development that drive T-cell responses from targets other than the S-protein when designing vaccines to combat coronaviridae infections.

Although T cells cannot prevent the initial entry of a virus into host cells, they can provide protection by recognizing viral peptides presented by human leukocyte antigens (HLAs) on the surface of host-infected cells, or antigen presenting cells (APCs). Several studies have demonstrated in SARS-CoV that virus specific CD8 T cells are required for mounting an effective immune response and viral clearance [9, 15-19]. A vaccine design that confers optimal protection may also need to involve the generation of memory T cell responses [20]. It has been shown that the activation of memory T cells specific for a conserved epitope shared by SARS-CoV and MERS-CoV is a potential strategy for developing coronavirus vaccines [21]. In addition, levels of memory T cell responses to SARS-CoV against peptides from its structural proteins were detected in a proportion of SARS-recovered patients, several years after infection [22, 23]. However, an adequate T cell response in isolation may not be sufficient. In a cohort study of 128 recovered SARS-CoV patients, the immune correlates of protection were investigated and broad CD8, CD4 and neutralizing antibody response were all shown to contribute to protection [24]. The CD4 T cell responses mainly clustered in the S-protein, presumably as B cell antibody responses to the S-protein requires the help of CD4 T cells specific to the same protein [25]. Given that in the before mentioned study by Mitchison et al that neutralizing antibody responses correlated with CD8 T cell responses against a broad set of CD8 T cell epitopes in the S-protein, a vaccine design that centers on the S-protein or any other viral protein will need to stimulate a broad CD8 response [26]. In the previous study, robust T cell responses correlated significantly with higher neutralizing antibody activity, consistent with the hypothesis that T cells play an important role in the generation of antibody responses in recovered SARS-CoV patients[24]. The concept of considering integrative CD8/CD4, and antibody immune parameters when designing a vaccine is also reinforced in an influenza mouse study which demonstrated that a universal T cell vaccine against Flu in the absence of a protective antibody response could result in a detrimental immunopathology in mice[27]. The importance of an integrative CD8, CD4 and B cell response in mounting a successful immune response against the present SARS-CoV-2 threat was well established at an early stage [28]. Activated CD4 T cells and CD8 T cells, in concert with antibodies against SARS-CoV-2 were recruited in a successful fight against the virus and recovery in an infected patient [28].

Many of the previous SARS-CoV studies have found promising CD8 targets [9, 15, 18, 24], including sustainable memory T cell responses [9, 15-17, 20-23] that recogise epitopes in proteins across the entire spectrum of the virus, although the S-protein have been reported to be enriched for dominant CD8 T cell responses [24]. Taken together, this supports the approach taken in this study, which is to map computationally, a broad epitope landscape across the global viral SARS-CoV-2 proteome, which includes integrated CD8, CD4 and B cell targets in the modeling. There has been some preliminary efforts submitted into preprint servers recently that describe epitope maps generated [29-31], however it appears that the emphasis in those approaches were based mostly on HLA binding. It is important to profile in whole viral proteome epitope screens, as carried out in this study using an extensive artificial intelligence platform, not only the candidates that may bind to the HLA molecule but also those CD8 epitopes that are naturally processed by the cell’s antigen processing machinery, and presented on the surface of the infected host cells. Layered on top of the antigen presentation predictions in the host infected cells, we also make predictions across the entire viral proteome that measure the likelihood that the peptides presented on the host infected cells are capable of being recognized by T cells that are not yet tolerized or deleted from a patient’s T cell repertoire. The subsequent immunogenic landscape of the SARS-CoV-2 that we present here is taken further to analyze the immunogenicity of all the non-synonymous variations across approximately 3400 different SARS-CoV-2 sequences, to map the trajectory of differential immunogenic potential between all the currently sequenced viral strains.

Any viable vaccine to tackle SARS-CoV-2 that incorporates T cell epitopes in its design would need to contain a constellation of overlapping epitopes that protect the vast majority of the human HLA population against the virus. In this study we attempt to demonstrate that the SARS-CoV-2 immunogenic landscape clusters into distinct groups across the spectrum of HLA alleles in the human population. Our predicted immunogenic landscape of the SARS-CoV-2 virus is therefore then processed through a robust comprehensive statistical Monte Carlo simulation, incorporating the integrative immune parameters, to identify epitope hotspots for a broad adaptive immune response across the most common HLA genetic makeup in the human population. The resulting epitope hotspots we identified may represent areas in the viral proteome that are most likely to be viable vaccine targets and represent blueprints for vaccine designs that may be universal in nature.

In order to rank-prioritize these potential universal epitope hotspots, and the peptides that underlie them at high resolution, the baseline peptide predictions are then taken through a graph based “digital twin” type simulation[32], to prioritize hotspots and the specific overlapping peptides that they comprise at a patient specific and population specific level. In addition, epitope hotspots containing viral epitopes that had high similarity with human peptides, especially those expressed in critical organs were removed from the blueprints. In this context, the digital twin information is the precise HLA genotype of an individual, and many virtual individuals are considered within a given population being analyzed. The HLA genotype is a key determinant of the immune response that a specific individual can mount against an infection, and it is also an important factor for determining whether a vaccine is effective in establishing immunity for the specific individual and a broader population (consisting of multiple diverse individuals). The candidate sequence targets that emerge from this computational analysis represent blueprints for potential vaccine designs modeled across the global human population

## Results

### The immunogenic landscape of SARS-CoV-2 reveals diversity among the different HLA groups in the human population

We carried out an epitope mapping of the entire SARS-CoV-2 virus proteome using cell-surface antigen presentation and immunogenicity predictors from the *NEC Immune Profiler* suite of tools. Antigen presentation (AP) was predicted from a machine-learning model that integrates in an ensemble machine learning layer information from several HLA binding predictors (in the case three distinct HLA binding predictors trained on ic50nm binding affinity data) and 13 different predictors of antigen processing (all trained on mass spectrometry data). The outputted AP score ranges from 0 to 1, and was used as input to compute immune presentation (IP) across the epitope map. The IP score penalizes those presented peptides that have degrees of “similarity to human” when compared against the human proteome, and awards peptides that are less similar. The resulting IP score represents those HLA presented peptides that are likely to be recognized by circulating T-cells in the periphery i.e. T-cells that have not been deleted or tolerized, and therefore most likely to be immunogenic. Both the AP and the IP epitope predictions are “pan” HLA or HLA-agnostic and can be carried out for any allele in the human population, however for the purpose of this study we limited the analysis to the 100 of the most frequent HLA-A, HLA-B and HLA-DR alleles in the human population. Class II HLA binding predictions were also incorporated into the large scale epitope screen from the IEDB consensus of tools [33], and B cell epitope predictions were performed using BepiPred [33]. The resulting epitope maps allowed for the identification of regions in the viral proteome that are most likely to be presented by host-infected cells using the most frequent HLA-A, HLA-B and HLA-DR alleles in the global human population. Epitope maps were created for all of the viral proteins and an example based on the IP scores for the S-protein is depicted in Figure 1A and for AP in Figure 1B, and illustrates distinct regions of the S-protein that contain candidate CD8 and CD4 epitopes for the 100 most frequent human HLA-A, HLA-B and HLA-DR alleles. Interestingly, the predicted B cell epitopes often map to regions of the protein that contain a high density of predicted T cell epitopes, thus the heat maps provide an overview of the most relevant regions of the SARS-CoV-2 virus that could be used to develop a vaccine. It is clear from Figure 1 that different HLA alleles have different Class I AP (and IP), and Class II binding properties. This strongly suggests, as one might anticipate, that the SARS-CoV-2 antigen presentation landscape (and IP) clusters into distinct population groups across the spectrum of different human HLA alleles. This trend is further illustrated in the hierarchichal-clustering map presented Figure 2 after the AP scores have been binarized. Figure 2 clearly demonstrates that some allelic clusters present many viral targets to the human immune system, while others only present a few targets, and some are unable to present any. This implies that different groups in the human population with different HLA’s will respond differentially to a T cell driven vaccine composed of viral peptides. Therefore in order to design the optimal vaccine that leverages the benefits of T cell immunity across a broad human population we need to predict “epitope hotspots” in viral proteome. These hotspots are regions of the virus that are enriched for overlapping epitopes, and/or epitopes in close spatial proximity, that can be recognized by multiple HLA types across the human population.

**Figure 1:**
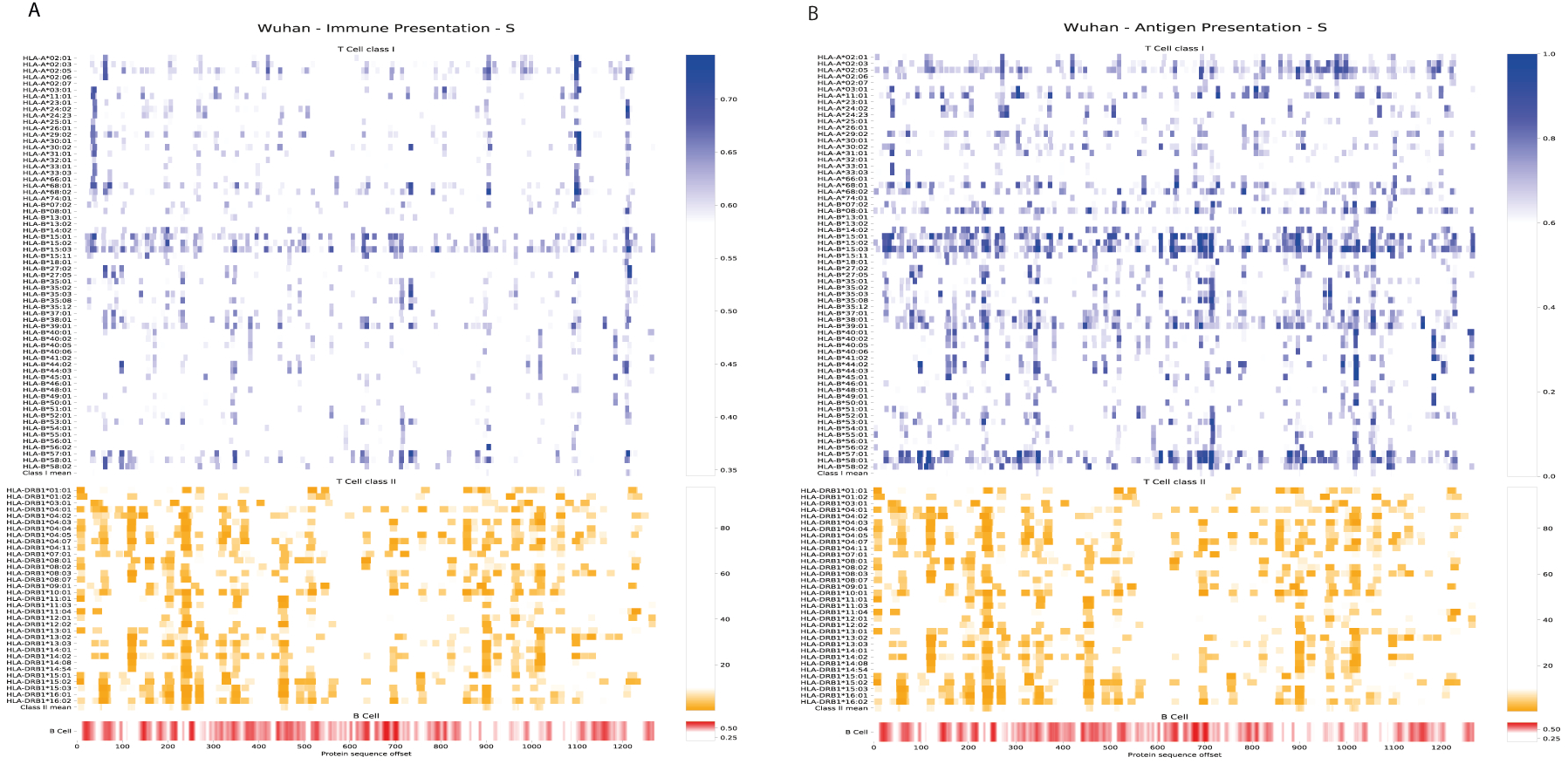
Epitope map of the S-Protein across the most frequent HLA-A. HLA-B and HLA-ORB alleles in the human population, Data is transformed such that a positive results for CD8 relates to 0.7 or above, and 0.1 or below for Class II, Broad coverage for CD8 and CD4 is demonstrated with overlaying Bcell antibody support.

**Figure 2:**
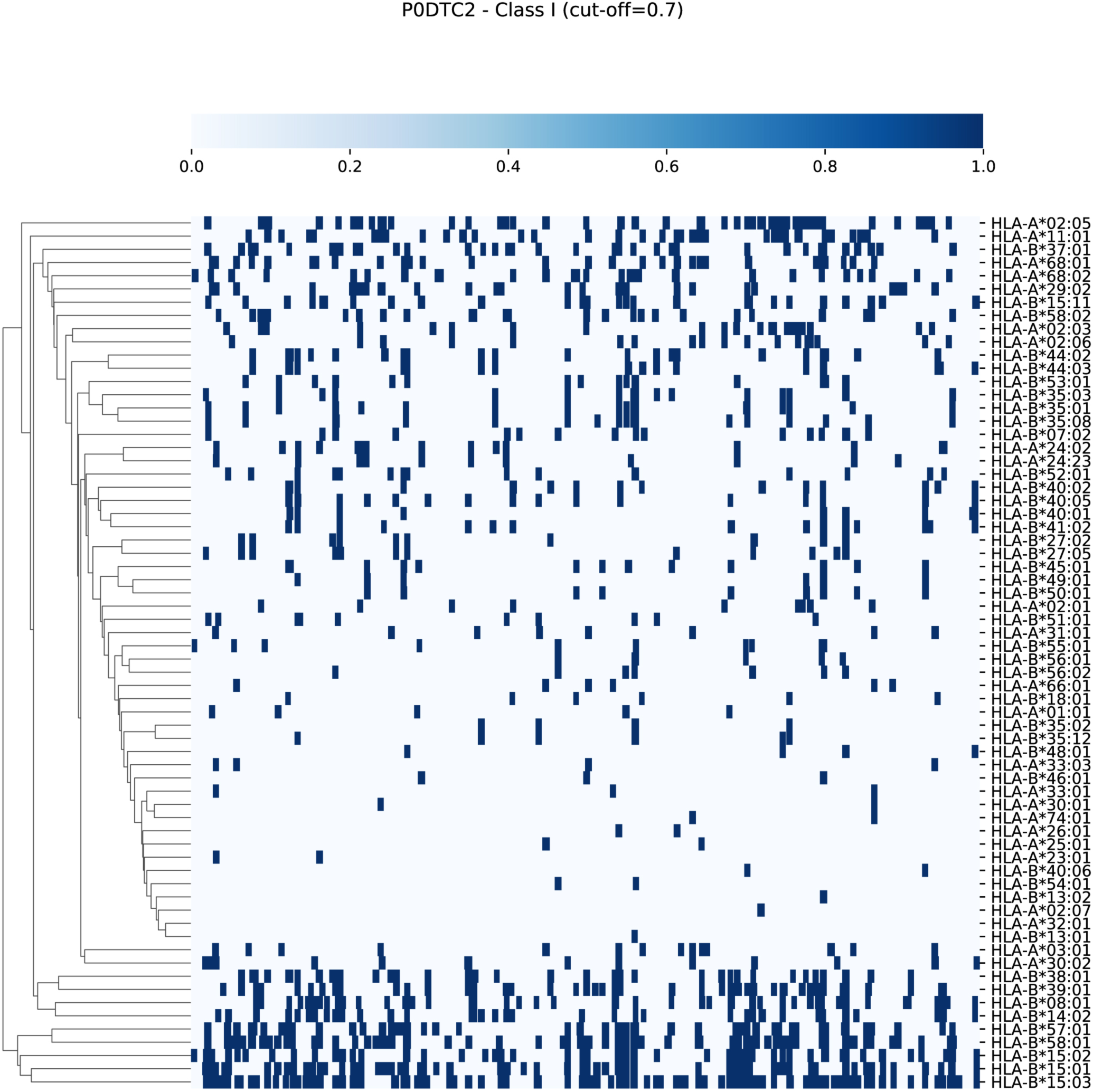
Hierarchical clustering of binary transformation of the epitope maps. Illustrated here for Class ICD8 epitopes in HLA-A and HLA-B alleles. Predictions for Class I are greater than 0.7 are transformed to 1. Illustrated for the S-Protein here for demon-stration purposes.

Prior to applying the *NEC Immune Profiler* suite of tools to map the SARS-CoV-2 viral proteome, it was important to first validate, to the extent that is possible from the limited number of validated SARS-CoV viral epitopes, that the T cell based AP and IP scores are predicting viable targets. We identified class I epitopes from the original SARS-CoV virus (that first emerged in the Guangdong province in China in 2002) that shared ≥90% sequence identity with the current SARS-CoV-2. Unfortunately, many of the published epitopes were identified using ELISPOT on PBMCs from convalescent patients and/or healthy donors (or humanised mouse models) where the restricting HLA was not explicitly deconvoluted. In order to circumvent this problem, we identified a subset of 5 epitopes where the minimal epitopes and HLA restriction had been identified using tetramers [6]. Four out of the 5 epitopes tested were identified as positive *i.e*. had an IP score of above 0.5 (see Table 1) demonstrating an accuracy of 80%. Although this was a very small test dataset, this provides us some degree of confidence that the *NEC Immune Profiler* prediction pipeline can accurately identify good immunogenic candidates and that the epitope hotspots identified by this analysis and subsequent analyses represent interesting targets for vaccine development.

**Table 1.**
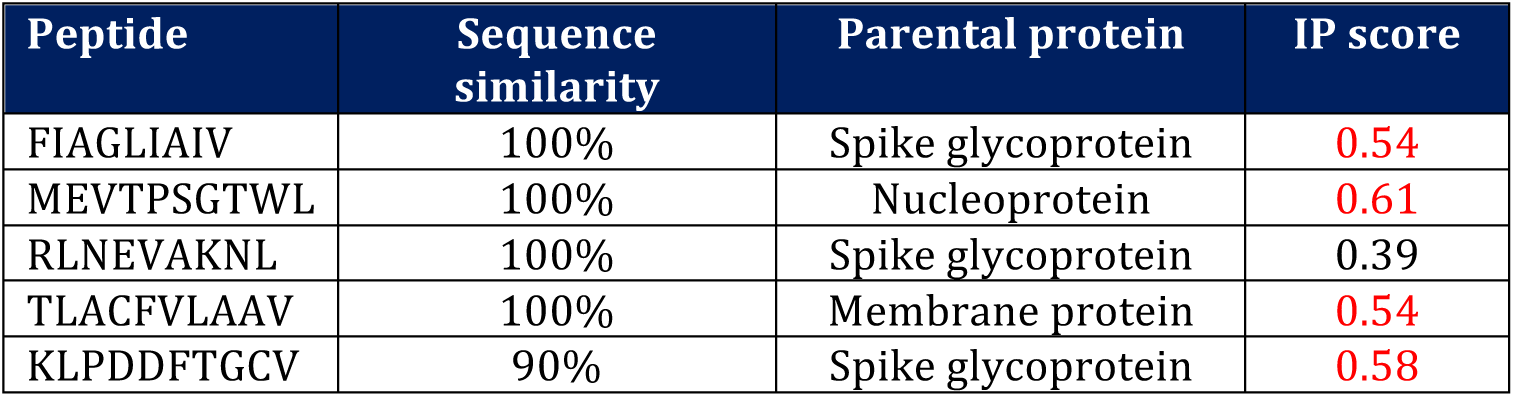

### A robust statistical analysis identifies epitope hotspots for a broad T cell response

In order to identify epitope hotspots that have the potential to be viable immunogenic targets for the vast majority of the human population, we first carried out a Monte Carlo random sampling procedure, on the epitope maps generated previously (for the Wuhan reference sequence exemplified in Figure 1 for the S-protein), to identify specific areas of the SARS-CoV-2 proteome that have the highest probability of being epitope hotspots (See Material and Methods). Three bin sizes were investigated for potential epitope hotspots; 27, 50 and 100. A statistic was calculated for each defined subset region of the protein (bin) from the set of 100 HLAs. The Monte Carlo simulation method was then used to estimate the p-values for each bin, whereby each bin represented a candidate epitope hotspot. The statistically significant bins that emerged from the simulation represented epitope hotspot or regions of interest for each protein analyzed. Epitope hotspots are built on the individual epitope scores, epitope lengths, and for each amino acid that they comprise. These scores are generated for each amino acid in the hotspots for all of the 100 HLA alleles most frequent in the human population. Based on the Monte Carlo analysis, the significant hotspots are those below a 5% false discovery rate (FDR), and represent regions that are most likely to contain viable T cell driven vaccine targets that can be recognized by multiple HLA types across the human population. A summary of the epitope hotspots identified across the entire spectrum of the virus is depicted in Figure 3 and reveals that the most immunogenic regions of the virus, that target the most frequent Human HLA alleles in the global population, are found in several of the viral proteins above and beyond the antibody exposed structural proteins, such as the S-protein.

**Figure 3:**
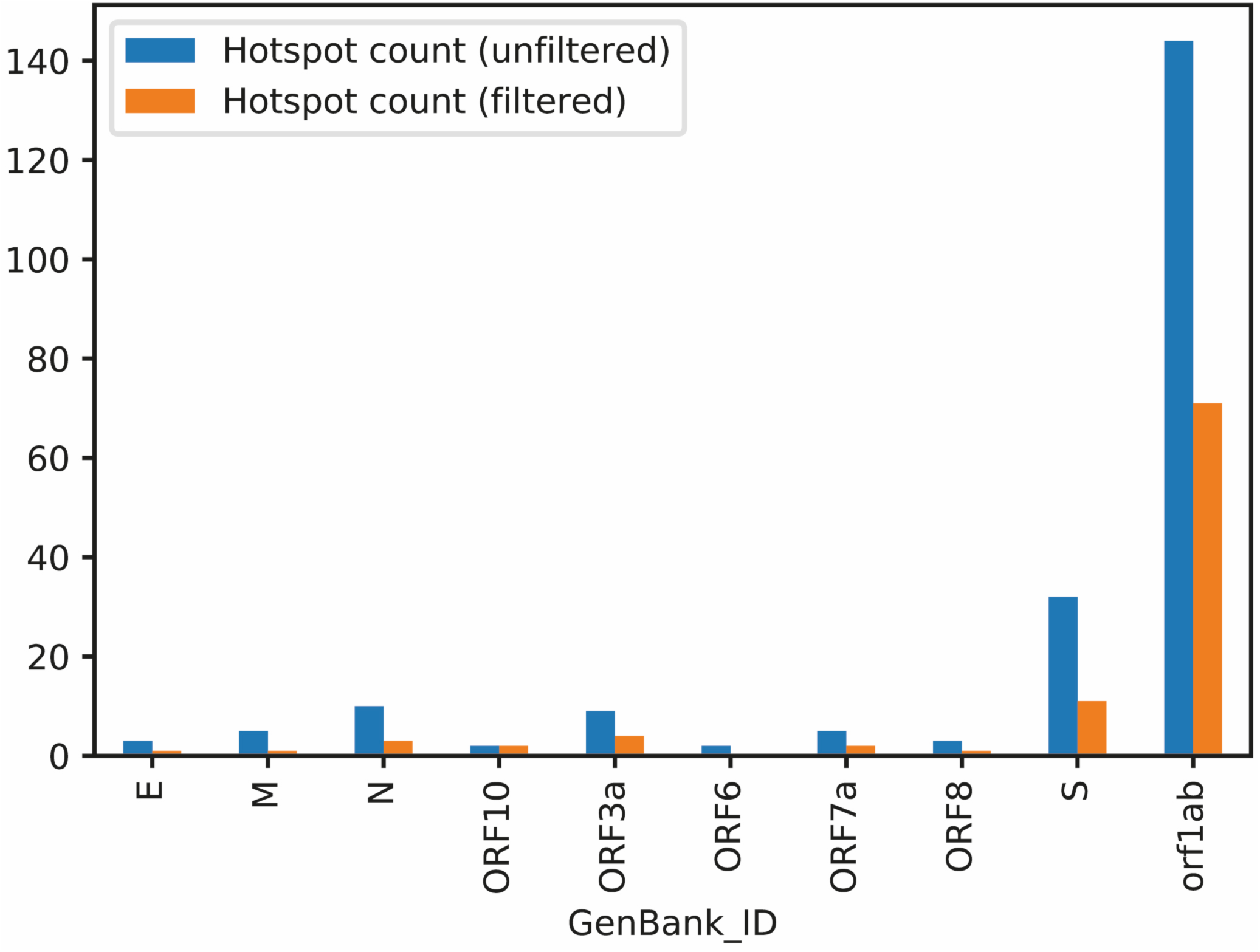
Epitope hotspots from the Monte Carlo analysis are captured across the entire viral proteome using filtering procedures for conserved and human self-peptides. The most abundant signal for hotspots is in the orf1 ab polyprotein.

### Conservation analysis identifies robust epitope hotspots in SARS-CoV-2

Although an early report in a preprint article demonstrated in a few sequences that the SARS-COV-2 genome has a lower mutation rate and genetic diversity compared to that of SARS-COV [34], another preprint study has demonstrated that there are evolving genetic patterns emerging in different strains of SARS-COV-2 in diverse geographic locations [35]. A universal vaccine blueprint should ideally also be able to protect populations against different emerging clades of the SARS-COV-2 virus and we therefore compared the AP potential of approximately 3400 virus sequences in the GISAID database against the AP potential of the Wuhan Genbank reference sequence. The outcome of that comparison is illustrated in Figure 4, and hints at a trend whereby SARS-COV-2 mutations seem to reduce their potential to be presented and consequently detected by the host immune system. Similar trends have been observed in chronic infections such as HPV and HIV.

**Figure 4:**
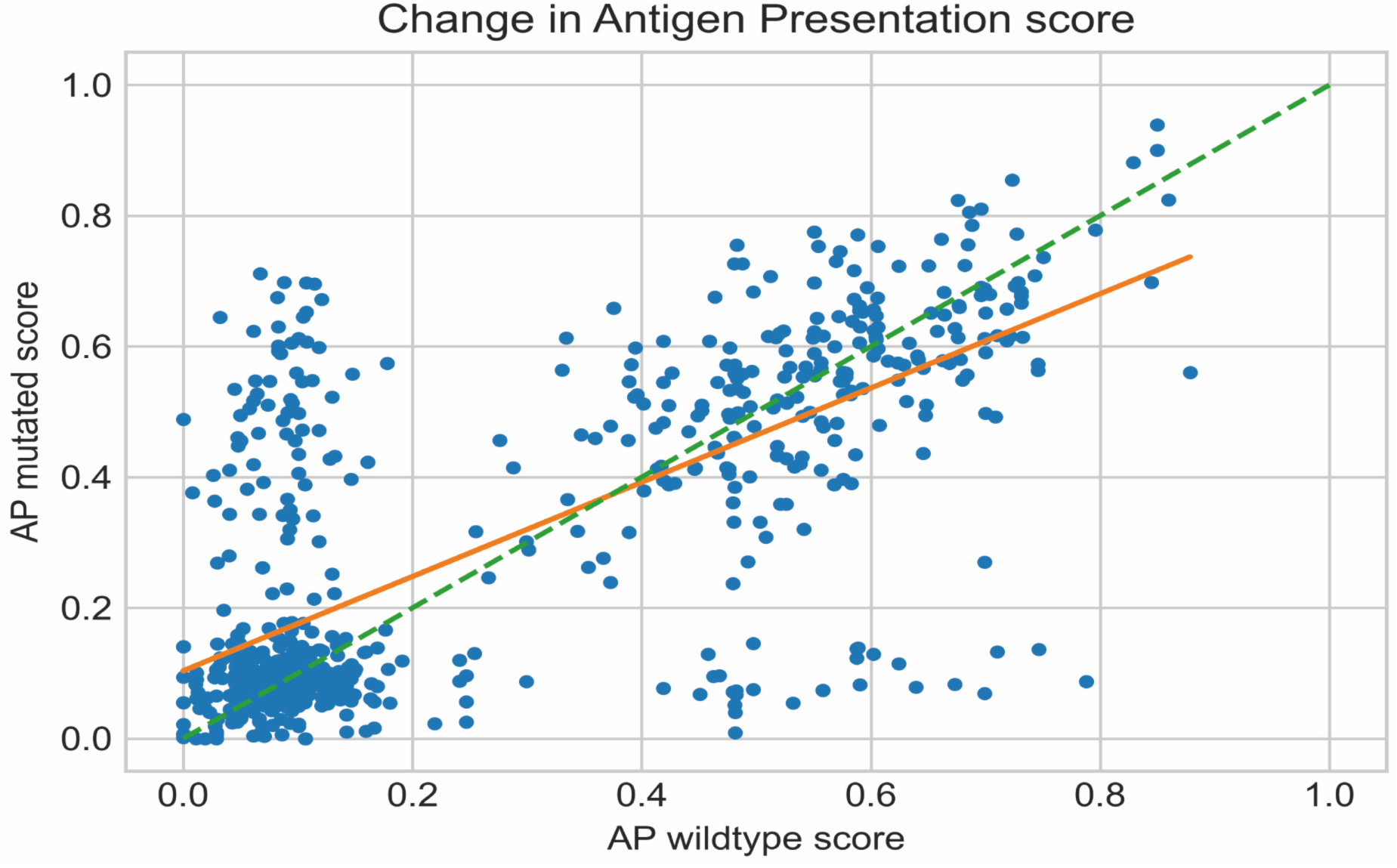
The scatter plot shows the mutated AP score (Y-axis) against its wildtype AP score (X-axis) for each protein variant. Each blue dot is one of the 1122 protein variants. The orange line is a least square fit with slope= 0.72. Variants on the green dashed line of slope 1 would show no change in AP after the introduction of the mutation.

In order to assess if these epitope hotspots are sufficiently robust across all the sequenced and mutating strains of SARS-CoV-2, we next used the epitope hotspot Monte Carlo statistical framework, and analyzed 10 sequences of the virus from among the 10 most mutated viral sequences from different geographical regions [36]. The vast majority of the hotspots were present in all of the sequenced viruses, however occasionally hotspots were eliminated and/or new hotspots emerged in these divergent strains as shown in Figure 5.

**Figure 5.**
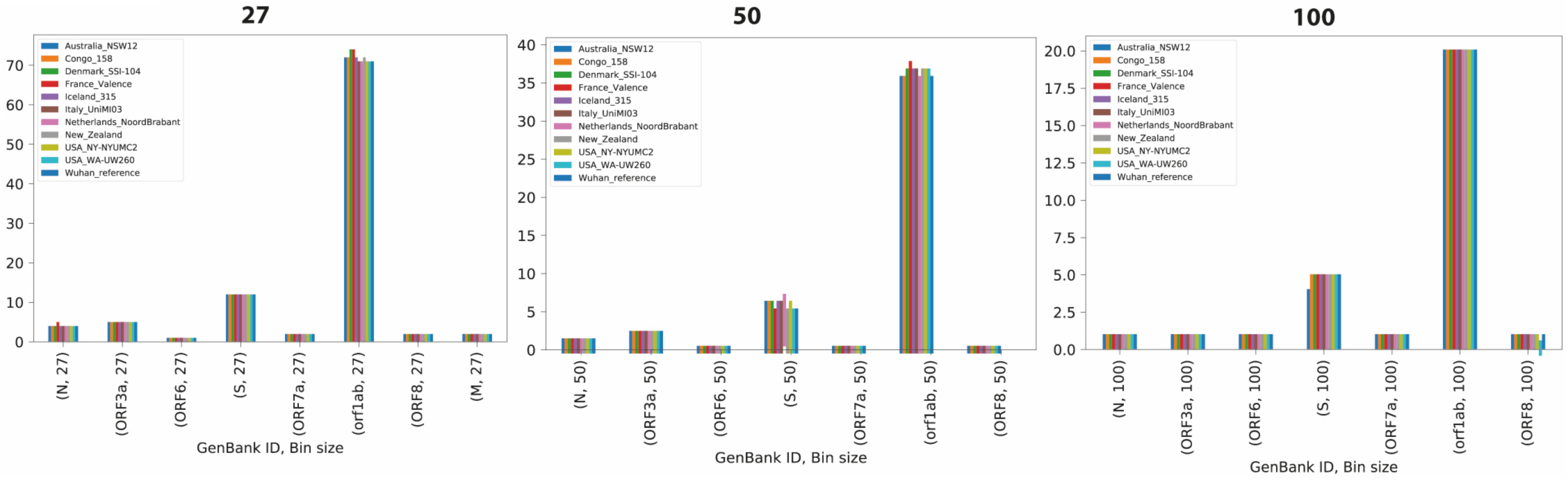
Application of the Monte Carlo epitope hotspot prediction method to 10 mutating virus sequences in diferent geographical locations. The hotspots for 10 mutated sequences compared to the Wuhan reference sequence are on the x- axis, the fequency of the epitope hotspots among the 11 total sequences. The frequencies are shown for 3 different epitope hotspot of bin lengths; 27 (left), 50 (centre) and 100 (right). It is clear that the epitope hotspots are robust across mutating sequences, while occasionally new epitope hotsptots emerge in some sequences in different geo-graphical locations.

Although the identified hotspots seem to be maintained across different viral strains, in order to design the most robust vaccine blueprint that will hopefully provide broad protection against new emerging clades of the SARS-COV-2 virus, the epitope hotspots were subject to a sequence conservation analysis. The goal of this analysis was to identify hotspots that appear to be less prone to mutation across thousands of viral sequences. We calculated a conservation score for each hotspot based on the consensus sequence of a protein (see Materials and Methods). Figure 6 shows conservation scores for the hotspots identified based on IP using different bin sizes. Only the epitope hotspots presenting a conservation score higher than the median conservation score were kept for further analysis. This allowed us to filter out a significant amount of less conserved epitope hotspots, that although have high immunogenic scores, harbor a higher degree of potential sequence variation. In addition, to reduce the potential for off-target autoimmune responses against host tissue we removed bins that contained exact sequence matches to proteins in the human proteome.

**Figure 6.**
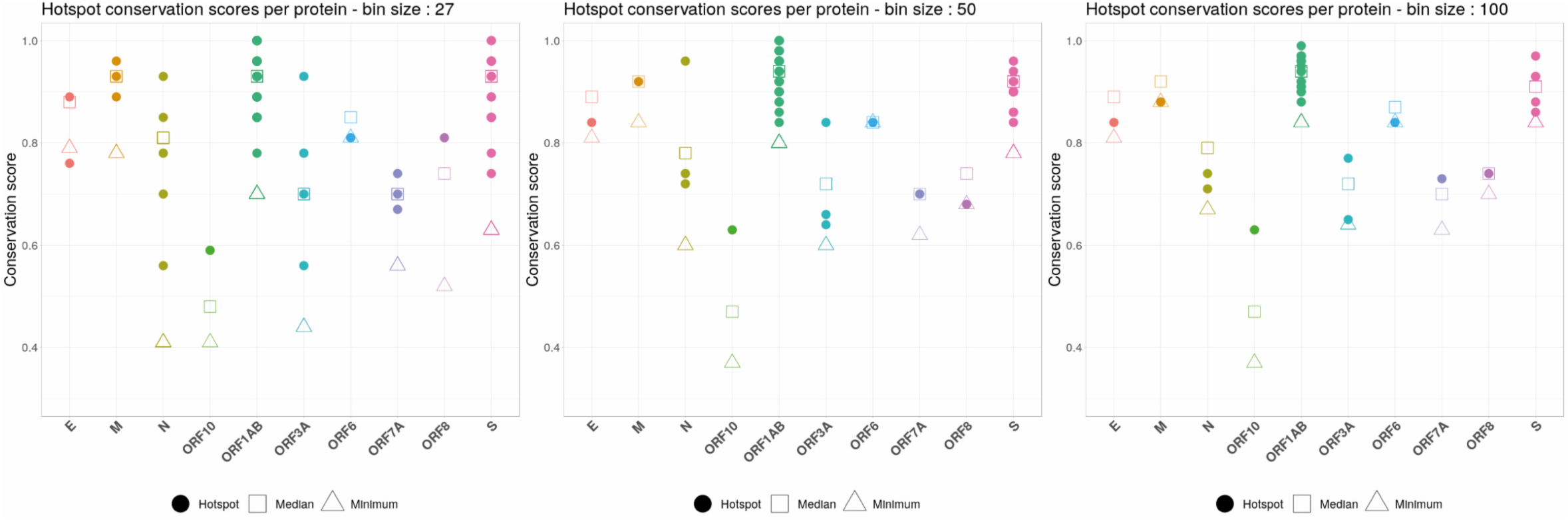
The scatter plots show the distribution of the hotspot conservation scores (Y-axis) for proteins in the viral genome (X-axis). Each plot contains the conservation scores of the hotspots identified based on immune presentation (IP), using different bin sizes: 27 (left), 50 (center), and 100 (right). A hotspot is represented by a filled coloured circle, while the median conservation score for a given protein is depicted by a hollow square. As a reference, upwards facing triangles show the minimum conservation score for that protein. Only hotspots with a conservation score above the median were taken for further considertion.

### A graph based “digital twin” optimization prioritizes epitopes hotspots to select universal blueprints for vaccine design

The Monte Carlo simulation identified well over 100 different hotspots of length 27, 50 or 100 amino acids, for both AP and IP. Even after filtering for conservation and self-similarity we were left with over 50 different hotpots for both the AP and IP based analyses. In order to develop a blueprint for viable universal vaccine against SARS-CoV-2, it is necessary to 1) cover with fidelity a broad proportion of the human population, and 2) prioritize the selection to even fewer regions (the exact number may depend on the size of the bin and the vaccine platform under consideration). Consequently, we need to identify the optimal constellation of hotspots, or relevant viral segments, that can provide broad coverage in the human population with a limited and targeted vaccine “payload”. In order to achieve this aim, we developed and applied (see Materials and Methods) a “digital twin” method, which models the specific HLA background of different geographical populations and used the method to identify optimal clusters of immunogenic epitope hotspots that will induce immunity in the broad human population. A graph-based mathematical optimization approach is then used to select the optimal combination of immunogenic epitope hotspots that will induce immunity in the broad human population. The results of this analysis are shown in Figure 7. The output clearly identified a subset of hotspots that may be combined to stimulate a robust immune response in a broad global population. An example hotspot for the ORF3a100-150 region is provided in the supplementary data file, which shows the amino acid sequence and its component Class I and Class II epitopes.

**Figure 7:**
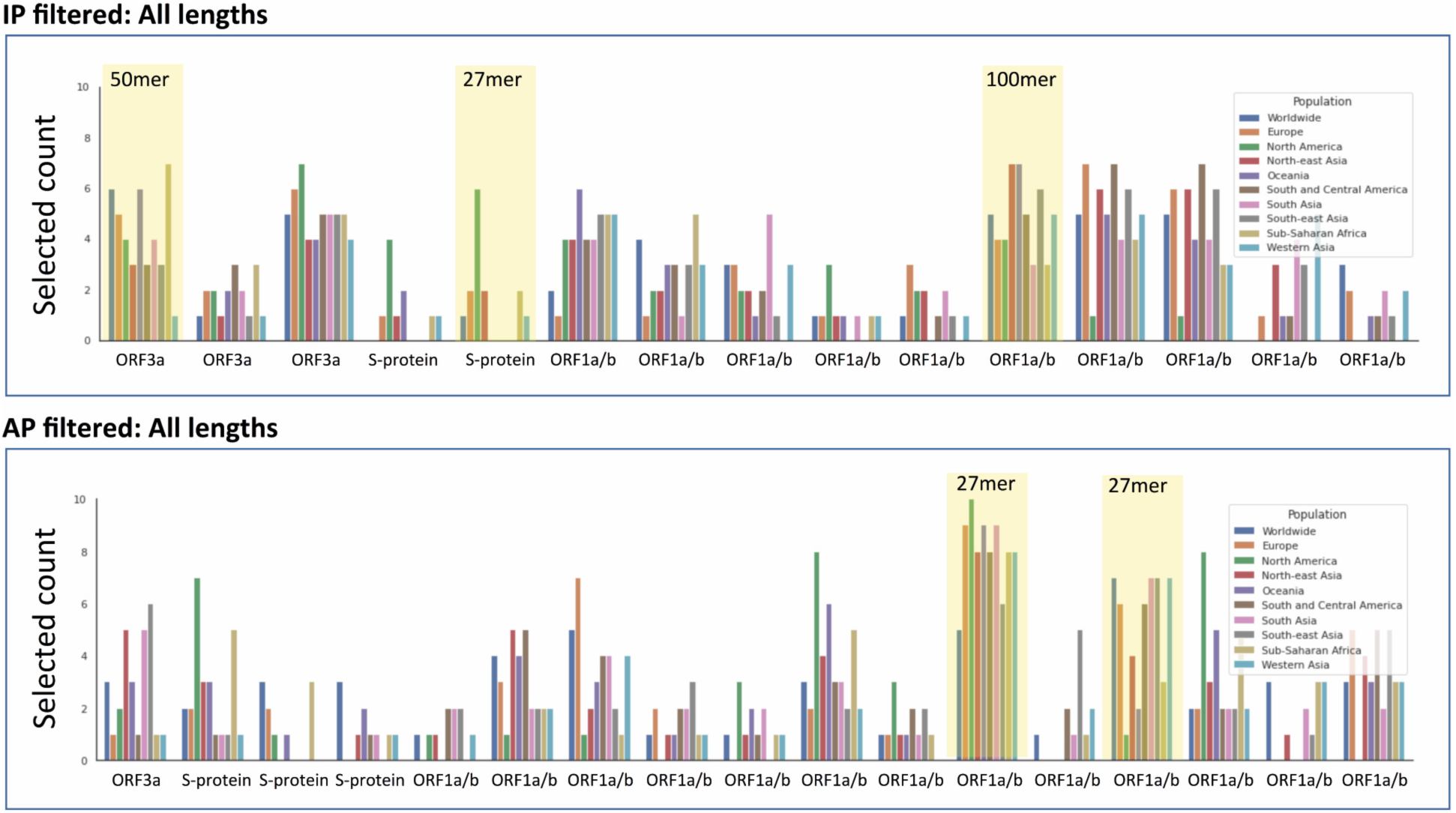
The plot illustrates a set of digital twin simulation experiments to identify effective epitope hotspots based on immune presentation (IP) and antigen presentation (AP), which broadly cover the population. The aim of the analysis was to select an optimum set of hotspots (respecting a given budget) such that the likelihood that each citizen (in a given modelled population) has a positive response is maximized (or, equivalently, that the log likelihood of no response for each citizen is minimized). Ten simulations were run for each region illustrated, where each simulation consisted of 10000 digital twins (see supplementary file for more details). This plot shows the number of times each hotspot was selected for use in one of the simulations. Each hotspot selected at least 10 times is shown on the x-axis, and the y-axis shows the selected counts per region. Each bar corresponds to a different region-specific simulation setting. A subset of 5 hotspots were selected based on their profile across the AP and IP digital twin analyses, which could theoretically provide coverage of >90% across a global population (highlighted in yellow).

## Conclusions

In order to effectively combat the SARS-CoV-2 pandemic a vaccine will need to protect the vast majority of the human population, and stimulate diverse T cell responses, against multiple viral targets including but not limited to the S-protein. To help achieve this ambitious aim, we have profiled the entire SARS-CoV-2 proteome across the most frequent 100 HLA-A, HLA-B and HLA-DR alleles in the human population and generated comprehensive epitope maps. We subsequently used these epitope maps as the basis for modeling the specific genetic HLA background of individual persons in a diverse set of different human populations using the most significant CD8 and CD4 T cell “epitope hotspots” in the virus. To the best of our knowledge this is the first computational approach that generates comprehensive vaccine design blueprints from large-scale epitope maps of SARS-CoV-2, in a manner that optimizes for diverse T cell immune responses across the global population. Underlying this approach are two novel methods that when integrated together result in a solution that is uniquely suited to achieving the objective of the study i.e. designing blueprints for universal vaccines. Firstly, a framework that leverages Monte Carlo simulations was developed to identify statistically significant epitope hotspot regions in the virus that are most likely to be immunogenic across a broad spectrum of HLA types. Secondly, a novel person-specific or “digital twin” type simulation using the actual HLA genotypes of approximately 22, 000 individuals prioritizes these epitope hotspots, to identify the optimal constellation of vaccine hotspots in the SARS-CoV-2 proteome that are most likely to promote a robust T cell immune response in the global population.

Importantly, the CD8 epitope maps that underlie these optimized epitope hotspots are based on our AP predictions of peptides presented on the surface of host-infected cells, and visible to the host’s CD8 T cells. Additionally, these antigen presented peptides are subject to our IP predictions that infer those specific epitopes that are most likely to activate a T cell in a host’s repertoire that has not been deleted or tolerized. These features confer unique properties to the epitope maps that underlie our epitope hotspot predictions and digital twin optimization. These properties differ from the SARS-CoV-2 epitope maps that have been reported in recent preprints since the outbreak of this virus, which mainly utilize predictions based on HLA binding [29-31].

A genomic analysis of approximately 3400 SARS-COV-2 sequences revealed that the epitope hotspots that we predict are robust across different evolving clades of the virus, which may be important in the design of universal vaccine blueprints. However, on average, mutations in the virus that cause amino acid changes in peptides seem to reduce their potential to be presented on the cell surface and consequently detected by the host immune system. We therefore apply filters on the vaccine blueprints that discard less sequence conserved hotspots, and hotspots that harbor peptides that have an exact match in the human proteome, before performing our digital twin simulation.

These results highlight the potential of looking beyond the S-protein and mining the whole viral proteome in order to identify optimal constellations of epitopes that can be used to develop efficacious and universal T-cell vaccines. The novel integrated methodological approaches described in this study may result in the design of diverse T cell driven vaccines that may help combat the SARS-CoV-2 pandemic, and bring much needed relief to the suffering global human population.

## Materials and Methods

### Generation of global epitope maps and amino acid scores

For a given HLA allele, the score allocated to an amino acid corresponds to the best score obtained by an epitope prediction overlapping with this amino acid. For Class I HLA alleles, the epitope lengths are 8, 9, 10 and 11, and predicted for antigen presentation (AP) or immune presentation (IP) of the viral peptide to host-infected cell surface, generated using the NEC Immune Profiler software. These Class I scores range between 0 and 1, where by 1 is the best score i.e., higher likelihood of being naturally presented on the cell surface (AP) or being recognized by a T-cell (IP). For class II HLA alleles, the only peptide length we have made predictions on is 15mers. The Class II were predictions were percentile rank binding affinity scores (not antigen presentation), so the lower scores are best (the scores range from 0 to 100, with 0 being the best score).

### Statistical framework for the detection of epitope hotspot epitope regions in different HLA populations

#### Input data

The data sets inputted into the statistical framework are epitope maps generated for each amino-acid position in all the proteins in the SARS-CoV-2 proteome, for all of the studied 100 HLA alleles. A score for any given amino acid was determined as the maximum AP or IP score that a peptide overlapping that amino acid holds in the epitope map. All peptide lengths of size 8-11 amino acids for class I, and 15 for class II were processed, generating one HLA dataset per viral protein. Each row in the dataset represents the amino acid epitope scores predicted for one HLA type.

#### Statistical framework

The central question that the statistical framework attempts to answer is: “are specific regions in a given viral protein enriched with higher immunogenic scores, with respect to a given set of HLA types, more than expected by chance?” To answer the question we implemented a hypothesis-testing framework inspired by work done in statistical genomics [37, 38].

#### HLA tracks

The raw input datasets are first transformed into binary tracks. For each class I HLA dataset, the epitope scores are transformed to binary (0 and 1) values, such that amino-acid positions with predicted epitope scores larger than 0.7 (for AP) and larger than 0.5 (for IP) are assigned the value 1 (positively predicted epitope), and the rest are assigned the value 0. Similarly, for class II HLA datasets, amino-acid positions with predicted epitope scores smaller than 10 are assigned the value 1, otherwise 0. These thresholds were relatively conservative. Each binary track can effectively be presented as a list of intervals of consecutive ones - segments, with consecutive zeros in between, forming inter-segments or gaps.

#### Test statistic

For a group of *k* HLA binary tracks, a test statistic *S*_*i*_ is calculated for each bin *b*_*i*_ of given size m, dividing the protein in n bins (e.g. m=100 amino-acids for the larger proteins). For a single HLA track, a test statistic *S*_*i*_ is calculated for each bin *b*_*i*_

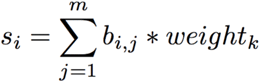

where the weight is by default 1.0, however can also represent frequency of the HLA track in the population under analysis.

Then, for *i=1..n*,

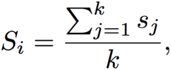

which is the average number of amino-acids predicted to be epitopes (epitope enrichment) of the bin *b*_*i*_, across the selected HLA types.

#### Null model

As with genomic tracks [38], analytical approaches to estimate the statistical significance of multiple observed HLA tracks are computationally intractable. An effective alternative to this problem is Monte Carlo-based simulations. A null model is defined, as the generative model of the HLA tracks, if they were generated by chance. From the null model, through sampling, arises the null distribution of the test statistic *Si*. The null model must reflect the complexities behind the nature of the HLA tracks. Amino acids in one HLA track will always form consecutive groups of length at least 8 (smallest peptide size used in the prediction framework). Similarly, amino acids with low epitope scores will also cluster together.

#### P-value estimation

To sample from the null model, each of the *k* HLA tracks is divided in segments and gaps, which are then shuffled to produce a randomized HLA track. This is repeated 10000 times, to produce 10000 samples of *Si* statistic for each bin. For each bin, the p-value is estimated as the proportion of the samples that are equal or larger then the truly observed enrichment. Further, the generated p-values are adjusted for multiple testing with the Benjamini–Yekutieli procedure to control for a false discovery rate (FDR) of 0.05.

### Graph based optimization in digital twin simulations of the epitope hotspots

We consider a population as a set *C* of “digital twin” citizens *c*, and a vaccine as a set *V* of vaccine elements *v*. We denote the likelihood that all citizens have a positive response to a vaccine as *P*(*R* = + |*C, V*). Our goal is to design a vaccine, that is, select a set of vaccine elements, to maximize this probability:

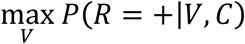

In this setting, maximizing the probability of positive response is the same as minimizing the probability of no response. Thus, we approach vaccine design by minimizing the probability of no response for the citizen who has the highest probability of no response *P*(*R* = − |*V, c*_*j*_):

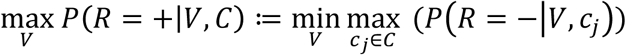

We consider that a vaccine causes a response if at least one of its elements causes a positive response. That is, the probability of no response is the joint likelihood that all elements fail. For a particular citizen *c*_*j*_, this probability is given as follows.

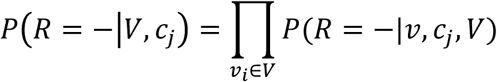

The original optimization problem can then be expressed as:

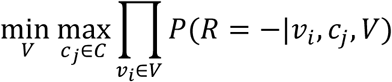

Since the logarithm function is monotonic, the value of *V* which minimizes the logarithm of the function also minimizes the original function.

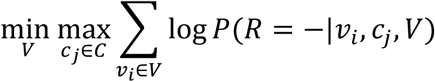

Further, we consider each citizen as a set of HLA alleles, and we assume that each vaccine element *v*_*i*_ may result in a response on each allele independently; we refer to the alleles for citizen *c*_*j*_ as *A*(*c*_*j*_). Thus, our final objective is as follows.

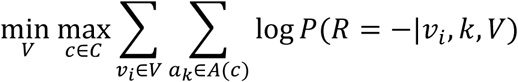

We approach this minimax problem as a type of network flow problem, with one set of nodes corresponding to vaccine elements, one set corresponding to HLA alleles, and one set corresponding to citizens. The goal is to select the set of vaccine elements such that the likelihood of no response is minimized for each citizen. Figure 8 gives an overview of the problem setting.

**Figure 8:**
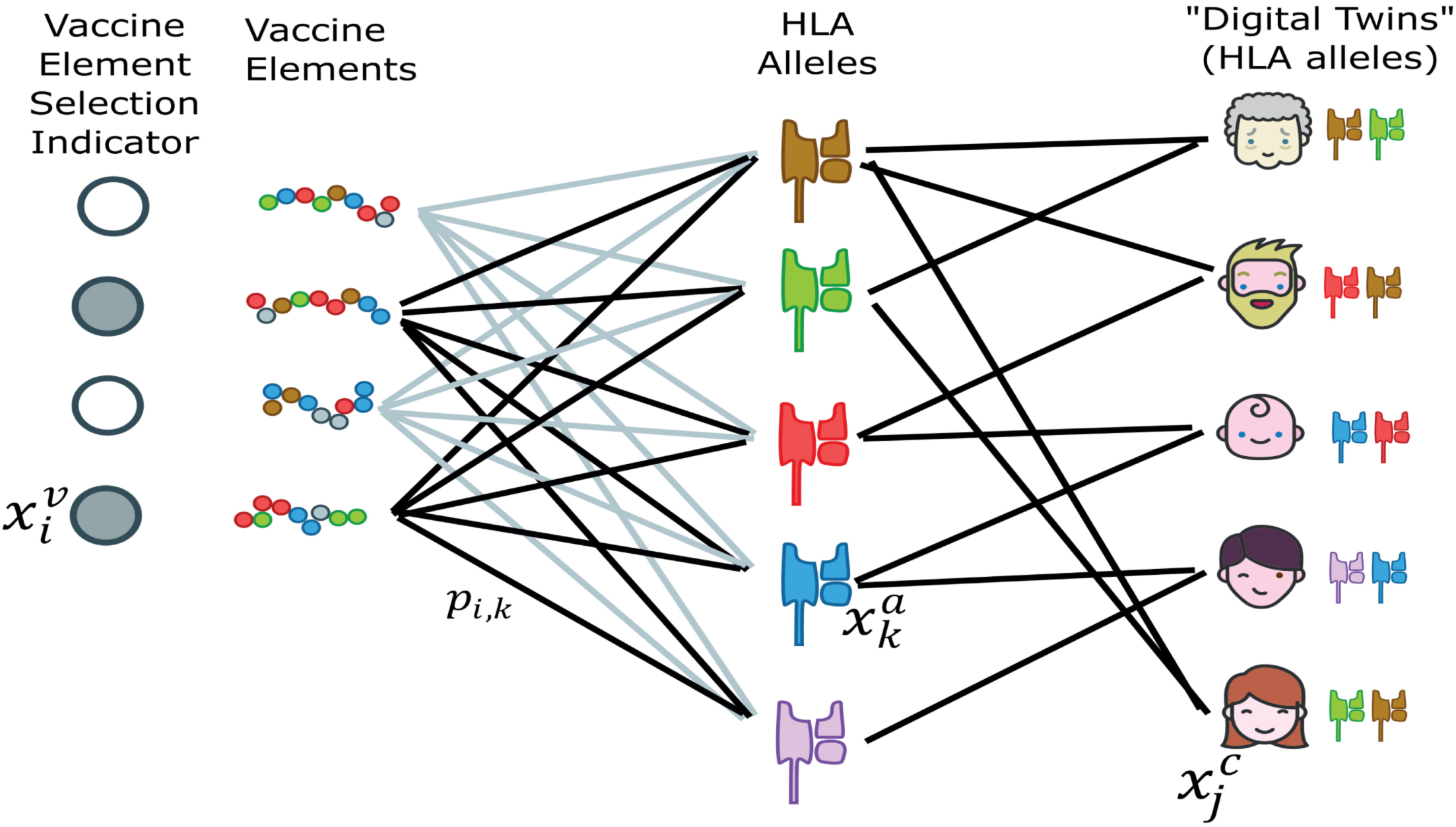
Schematic of the problem setting. The vaccines elements were the significant epitope hotspots that emerged from the statistical hotspot detection framework

### Vaccine design process

Concretely, we approach the vaccine design process in four steps:

1. Select a set of candidate vaccine elements for inclusion in the vaccine.
2. Create a set of “digital twin” citizens for a population of interest, where a digital twin is a set of HLA alleles.
3. Create a tripartite graph in which the nodes correspond to vaccine elements, HLA alleles, and citizens; edges correspond to relevant biological terms described below.
4. Select a set of vaccine elements (respecting a given budget) such that the likelihood that each citizen has a positive response is maximized (or, equivalently, that the log likelihood of no response for each citizen is minimized).

Each step is described in more detail in the supplementary methods

### Variant immunogenic potential across the mutating sequences of SARS-CoV-2

We downloaded all the strains available in the GISAID database [36] as of 31.03.2020, and ran them through the Nexstrain/Augur software suite with default parameters [39]. We parsed the resulting phylogenic tree to obtain all protein variants. For each we computed a wildtype score and a mutated Antigen Presentation (AP) score for HLA-A*02:01. The mutated score is the maximum AP score among the nine possible 9-mers peptides that include the variant. The wildtype score is the maximum AP score for the 9-mers at the same positions in the reference (Wuhan) strain.

### Epitope hotspot conservation scores

For each protein within the viral genome, the set of unique amino acid sequences was compiled from all the strains available in the GISAID database [36] as of 29.03.2020. These sets were individually processed using the Clustal Omega (v1.2.4) [40] software via the command line interface with default parameter settings. The software outputs a consensus sequence that contains conservation information for each amino acid within the protein sequence. As such, an amino acid depicted as an “*” at position *i* within the consensus sequence translates to that amino acid being conserved at position *i* among all the input sequences[40]

The hotspot offsets were then used to extract their respective consensus sub-sequence. For each hotspot, the conservation score was calculated as the ratio of “*” within its consensus sub-sequence to the total length of the sub-sequence. Accordingly, each hotspot was assigned a conservation score between 0 and 1, with 1 representing a perfect conservation across all available strains.

The median conservation score was calculated by sampling 1,000 sub-sequences equal to the hotspot size from the entire consensus sequence of a protein. Each sample was assigned a conservation score and the median value from all 1,000 conservation scores was calculated. The minimum conservation score was calculated using a sliding window approach, with the window size being equal to the hotspot size. For each increment, a conservation score was calculated and the resulting minimum conservation score was kept.

## Supplementary methods

### The digital twin simulation framework

#### Step 1. Select a set of candidate vaccine elements

Some of these candidate vaccine elements will be selected for inclusion in a vaccine. Two examples of vaccine elements are: (1) short peptide sequences, such as 9-mer amino acid sequences; (2) long peptide sequences, such as 27-mer amino acid sequence which may be based on a short peptide sequence and include flanking regions.

Each vaccine element *v*_*i*_ is associated with a cost 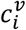, while a total budget *b* is available for including elements in the vaccine. The description of the budget and costs depend on the vaccine platform.

Some vaccine platforms are mainly restricted to a fixed number of vaccine elements; in this case, each cost 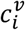 will be 1, and the budget will indicate the total number of elements which can be included.

Some other vaccine platforms are restricted to a maximum length of included elements. In this case, each cost 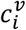 will be the length of the vaccine element, and the budget will indicate the maximum length of elements which can be included.

#### Step 2. Create a set of “digital twin” citizens

Our approach is based on simulating a set of “digital twin” citizens. In this work, we focus on vaccine elements whose effects are determined, in part, by the HLAs of each citizen. Thus, each digital twin corresponds to a set of HLA alleles.

It is known^1^ that citizens from different regions of the world tend to have different sets of HLA alleles; further, some combinations of HLA alleles are more common than others. We use full HLA genotypes from actual citizens available from high-quality samples in the Allele Frequency Net Database^1^ (AFND) to accurately model these relationships.

### Creating a distribution over genotypes for each region

In particular, AFND assigns each sample to a region based on where the sample came from (e.g., “Europe” or “Sub-Saharan Africa”). In a first step, we create a posterior distribution over genotypes in each region based on the observations and an uninformative (Jeffreys) prior distribution.

Specifically, we collect all genotypes observed at least once across all regions; we assign an index *g* to each genotype, and we call the total number of unique genotypes as *G*. Second, we specify a prior distribution over genotypes. We use a symmetric Dirichlet distribution with concentration parameter of 0.5 because this distribution is uninformative in an information theoretic sense and does not reflect strong prior beliefs that any particular genotypes are more likely to appear in any specific region. For each region, we then calculate a posterior distribution over genotypes as a Dirichlet distribution as follows.

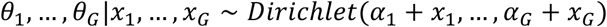

where *α*_*g*_ is the (prior) concentration parameter for the *g*^*th*^ genotype (always 0.5 here) and *x*_*g*_ is the number of times the *g*^*th*^ genotype was observed in the region.

We can now use this distribution to sample genotypes from a region using a two-step process.

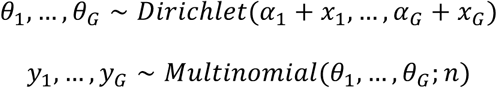

where *n* is the desired number of genotypes to sample from the region, and *y*_1,_ …, *y*_*G*_ are the counts of each genotype in the sample.

### Creating a set of “digital twin” citizens

We create a set of digital twin citizens using a two-step approach. Our method must be given the population size *p*, as well as a distribution over regions. Concretely, the input is a Dirichlet distribution over the regions, as well as *p*. (We note that this Dirichlet is completely independent of those over genotypes discussed in the previous section.) The number of citizens from each region is sampled using the same two-step sampling process described above.

Second, the genotypes for each region are sampled using the posterior distributions over genotypes discussed above.

#### Step 3. Create a tripartite graph

We next use the vaccine elements and digital twins to construct a tripartite graph that will form the basis of the optimization problem for vaccine design. The graph has three sets of nodes:

1. All candidate vaccine elements identified in Step 1
2. All HLA alleles in all digital twin genotypes
3. All digital twins

The graph also has two sets of weighted edges:

1. An edge from each vaccine element *v*_*i*_ to each HLA allele *a*_*k*_. The weight of this edge is log *P*(*R* = − | *v*_*i*,_ *a*_*k*_), that is, the likelihood of no response for the allele from that particular vaccine element. (We describe below an approach for calculating this value for short peptides.)
2. An edge from each allele to each citizen which has that allele in its genotype. The weight of these edges is always 1.

As an intuition, we call the edges from a vaccine element to an allele (and, then, from the allele to each patient with that allele) as “active” when the vaccine element is selected. Then, the log likelihood of response for a citizen is the sum of all active incoming edges. That is, the flow from selected vaccine elements to the citizens gives the likelihood of no response for that citizen.

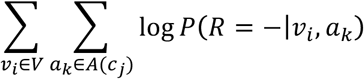

This definition does not include *V* in the conditioning set of the likelihood. Thus, it does not account for interactions among vaccine elements, such as immunodominance.

### Calculating the likelihood of no response for a given digital twin and vaccine elements

We now describe example approaches for calculating log *P*(*R* = −| *v*_*i*,_ *a*_*k*_) for three types of vaccine elements. Our vaccine design approach is applicable for any approach which assigns a value for log *P*(*R* = −| *v*_*i*,_ *a*_*k*_).

1. *Short peptide sequences.* Most short peptide prediction engines^1^ compute some sort of a score that a peptide will result in some immune response (e.g., binding, presentation, cytokine release, etc.), and this score generally takes into account a specific HLA allele. In some cases, this is already a probability, and in others, it can be converted into a probability using a transformation function, such as a logistic function. Thus, the prediction engines give *P*(*R* = +| *v*_*i*,_ *a*_*k*_), where *v*_*i*_ is the peptide and *a*_*k*_ is the allele. We then take log*P*(*R* = −| *v*_*i*,_ *a*_*k*_) = log[1 − *P*(*R* = +| *v*_*i*,_ *a*_*k*_)].
2. *Long peptide sequences*. Longer peptide sequences may include multiple short peptide sequences with different scores from the prediction engine. An example approach to calculate log *P*(*R* = −| *v*_*i*,_ *a*_*k*_), where *v* is the long peptide sequence, is to take the minimum (i.e., best) log *P*(*R* = −| *p, a*_*k*_), where *p* is any short peptide contained in *v*_*i*_.
3. *Longer amino acid sequences*. Longer amino acid sequences may contain even more short peptide sequences, and the same approach used for *long peptide sequences* can be used here.

#### Step 4. Selecting a set of vaccine elements

Finally, we pose the vaccine design problem as a type of network flow problem through the graph defined in Step 3. In particular, the minimization problem can be posed as an integer linear program (ILP); thus, it can be provably, optimally solved using conventional ILP solvers.

### Handling the minimax problem

As previously described, our goal is to choose the set of vaccine elements which minimize the log likelihood of no response for each patient.

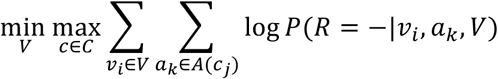

We ignore any interactions among vaccine elements, so the minimax problem simplifies as follows.

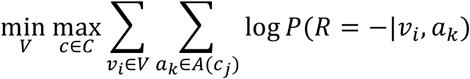

In practice, we can remove *V* from the conditioning set. Thus, the terms inside the summation are exactly those calculated in Step 3 as the weights on the edges in the graph.

Standard ILP solvers cannot directly solve this minimax problem; however, we use the standard approach of a set of surrogate variables to address this problem. In particular, we define 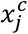 to be the log likelihood of no response for citizen *c*_*j*._ That is, 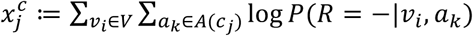. Further, we define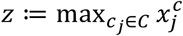; that is, *z* is the maximum log likelihood that any citizen does not respond to the vaccine (or, alternatively, the minimum log likelihood that any citizen will respond to the vaccine). Finally, then, our aim is to minimize *z*.

### ILP formulation

Our ILP formulation consists of three types of variables:

- 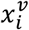: one binary indicator variable for each vaccine element which indicates whether it is included in the vaccine for the given population. We usually index vaccine elements with *i*.
- 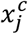: one continuous variable for each citizen in the population which gives the log likelihood of no response for that citizen. We always index citizens with *j*.
- 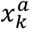: one continuous variable for each HLA allele which gives the log likelihood of no response for that allele. We always index alleles with *k*.
- *z*: one continuous variable which gives the maximum log likelihood that any citizen does not respond to the vaccine. (Our goal will be to minimize this value.)

Additionally, the ILP uses the following constants:

- *p_i,k_*: the log likelihood that vaccine element *v*_*i*_ does not cause a response for allele *k*.
- 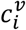: the “cost” of vaccine element *v*_*i*_.
- *b*: the maximum cost of vaccine elements which can be selected.

Finally, the ILP uses the following constraints:

- 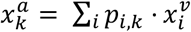: one constraint for each allele which gives the log likelihood that at least one selected peptide results in a positive response for that allele
- 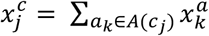: one constraint for each citizen which gives the log likelihood that at least one selected peptide results in a positive response for at least one allele for that citizen. (That is, this is the likelihood of a positive response for this citizen.)
- 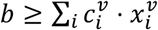: the vaccine elements we select cannot exceed the budget
- 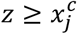: as discussed above, we use *z* as an approach to solve the minimax problem. These constraints imply that *z* is the minimum log likelihood that any individual patient will respond to the vaccine.

### Objective

The objective of the ILP is to minimize *z*.

The setting of the binary 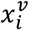 variables corresponds to the optimal choice of vaccine elements for the given population.

### Relationships to max-flow and other problems with provably efficient solutions

This is highly-related to a number of efficiently solvable network flow problems. Our problem is essentially a min-flow problem with multiple sinks, where each citizen is a sink; however, our aim is to minimize the flow to each individual sink rather than the flow to all sinks. In particular, rather than the “sum” operator typically used to transform multiple sink flow problems into a single-sink problem, we would need a (non-linear) “min” operator. Thus, efficient min-flow formulations are not applicable in this setting.

## Footnotes

1 Cao, K.; JillHollenbach; Shi, X.; Shi, W.; Chopek, M. & Fernández-Viña, M. A. Analysis of the frequencies of HLA-A, B, and C alleles and haplotypes in the five major ethnic groups of the United States reveals high levels of diversity in these loci and contrasting distribution patterns in these populations. Human Immunology, 2001, 62, 1009-1030.

2 http://www.allelefrequencies.net/

3 Jensen, K. K.; Andreatta, M.; Marcatili, P.; Buus, S.; Greenbaum, J. A.; Yan, Z.; Sette, A.; Peters, B. & Nielsen, M. Improved methods for predicting peptide binding affinity to MHC class II molecules. Immunology, 2018, 154, 394-406.

